# Two mutations P/L and Y/C in SARS-CoV-2 helicase domain exist together and influence helicase RNA binding

**DOI:** 10.1101/2020.05.14.095224

**Authors:** Feroza Begum, Arup Kumar Banerjee, Prem Prakash Tripathi, Upasana Ray

## Abstract

RNA helicases play pivotal role in RNA replication by catalysing the unwinding of complex RNA duplex structures into single strands in ATP/NTP dependent manner. SARS coronavirus 2 (SARS-CoV-2) is a single stranded positive sense RNA virus belonging to the family *Coronaviridae*. The viral RNA encodes non structural protein Nsp13 or the viral helicase protein that helps the viral RNA dependent RNA polymerase (RdRp) to execute RNA replication by unwinding the RNA duplexes. In this study we identified a novel mutation at position 541of the helicase where the tyrosine (Y) got substituted with cytosine (C). We found that Y541C is a destabilizing mutation increasing the molecular flexibility and leading to decreased affinity of helicase binding with RNA. Earlier we had reported a mutation P504L in the helicase protein for which had not performed RNA binding study. Here we report that P504L mutation leads to increased affinity of helicase RNA interaction. So, both these mutations have opposite effects on RNA binding. Moreover, we found a significant fraction of isolate population where both P504L and Y541C mutations were co-existing.

## Introduction

SARS-CoV2 or the novel coronavirus 2019 is a member of the family *Coronaviridae*. It is the seventh member of human coronaviruses of this family. There is no vaccine or antiviral against this virus till date.

Coronaviruses are single stranded positive sense RNA viruses. The viral RNA has 5’ cap and a 3’ polyA tail and encodes two different classes of proteins, the structural and the non-structural proteins [1]. The structural proteins participate in forming the structure of the virus particle whereas the non-structural proteins play crucial roles in various steps of the virus life cycle after entry into the host cells. After the virus enters the host cell, the virus undergoes uncoating and the viral genetic material is released in the host cell cytoplasm where the viral positive sense RNA undergoes translation to form the various viral proteins. The viral proteins are formed in the form of polyprotein that gets processed by host cell proteases and viral protease. Followed by synthesis of viral non-structural proteins the viral RNA undergoes replication. Replication is important as this gives rise to new viral RNA strands that get later packaged into progeny virus particles. The viral RNA replication is primarily aided by two proteins, the viral RNA dependent RNA polymerase (RdRp) and the viral helicase.

Helicase protein is important for executing viral RNA replication wherein this enzyme catalyzes unwinding of complex RNA structures/ duplex structures into single strand for RdRp to initiate RNA synthesis [1]. The unwinding of viral RNA is accomplished in an ATP dependent fashion. Since RNA helicase plays a very important role in viral life cycle it is also one of the major therapeutic targets.

In this study we have compared SARS-CoV-2 helicase protein sequences from different geographical locations to identify possible mutations that might have emerged and propagated in the population in significant fractions. Such variants that are significantly high in number and have managed to survive the selection pressure might have important role in virus adaptation and evolution. Thus, such mutant proteins need to be characterized in depth to understand the role of such mutations in the viral life cycle. These mutants could be potential therapeutic targets or handles for better therapeutic interventions. Or, identification of the mutation hot spots in such crucial proteins could help identify the conserved regions of the protein that could be targeted in a generic fashion.

## Methods

### Sequence source

1549 SARS-CoV-2 Nsp13/helicase sequences were downloaded from GenBank. Out of these 1361 sequences were from North America, 5 from South America, 139 from Asia and 44 from Europe. These numbers reflect the full sequences with high coverage and available in annotated forms in the GenBank. For the current study, YP_009725308.1 was used as the wild-type sequence/ reference.

### Sequence Alignments

The sequences with respect to SARS-CoV-2 helicase were aligned using multiple sequence alignment tool of CLUSTAL Omega in batches.

### Protein motif scan and homology modelling

Protein domain search and motif scan were performed using PROSITE (ExPASy Bioinformatics resource portal) and MOTIF search servers respectively. Homology modelling for the mutant helicases were performed using SWISS-MODEL homology-modelling pipeline’s structure assessment tool [2] using the existing SARS-CoV-2 helicase model [ID: 6jyt.2.pdb] as the wild-type template. Template was selected based on maximum sequence similarity with the mutant helicase sequences.

### Secondary structure

CFSSP (Chou and Fasman secondary structure prediction) server was used to predict the secondary structures of SARS-CoV2 wild-type and mutant helicases.

### Protein structure stability and RNA binding

mCSM NA was used to detect the effect of mutations on helicase-RNA binding. Structural stability was assessed using two different tools, SAAMBE 3D [3] and MUPro.

### Protein dynamics

To study the effect of mutation on the stability and flexibility of the helicase protein, the available SARS-CoV-2 helicase structure was uploaded on DynaMut software (University of Melbourne, Australia) [4]. Change in vibrational entropy; the atomic fluctuations and deformation energies due to mutation were determined.

## Results and Discussion

Function of RNA helicase is important in the viral RNA replication process as it unwinds the RNA duplex into single strands so that the RdRp can carry out RNA synthesis. Thus, helicase compliments the function of RdRp. Here, we studied 1549 sequences corresponding to SARS-CoV-2 RNA helicase from different geographical locations. Sequence alignments and comparison with reference sequence form China revealed two amino acid positions that were mutated in more than 400 samples in case of each type. These mutations accounted to nearly 27-30% of the total sequences analysed (Figure 1) although this number might further change with more sequences been checked. Both these mutations were found to co-exist in 409 sequences of the North American isolates. At position 504 of the helicase, proline (P) was replaced by Leucine (L) (mutation P504L) and at 541 position, tyrosine (Y) got mutated to Cysteine (C) (mutation Y541C). In our earlier study we had reported mutation at position 504 but had not characterized the same in depth with respect to location of mutation and effect on RNA binding [5]. To check the location of these mutations on the helicase protein, we subjected the sequence of SARS-CoV-2 sequence to motif scan (Figure 1C, D). We found that the mutation P504L was located at the second AAA domain (between residues 486 to 575). AAA family of domains have conserved modules which bring together active site elements of adjacent subunits for ATP hydrolysis [6]. This is required for generation of mechanical force for translocation of substrate which in case of SARS-CoV2 helicase should be the RNA duplex that needs to unwind for the replication to initiate or progress. Thus, a mutation in this domain could have modulatory function on RNA replication by possibly modulating either the efficiency of ATP hydrolysis or the dynamics/speed of RNA unwinding. In a report by Wei et al, it was shown that a P to L mutation in the ATPase p97 showed enhanced ATPase activity [6] and desensitized this protein against its inhibitors. In another study by Lin et al it was demonstrated that a mutation from P to L in the reverse transcriptase domain of Hepatitis B virus (HBV) increased replication competency of the enzyme [8]. Although in our study the location of mutation and the enzyme are different than those of cited studies, it supports the likelihood that P504L mutation might lead to better helicase activity and thus might in turn enhance or expedite RdRp activity by better RNA unwinding and translocation processes.

**Figure 1.**
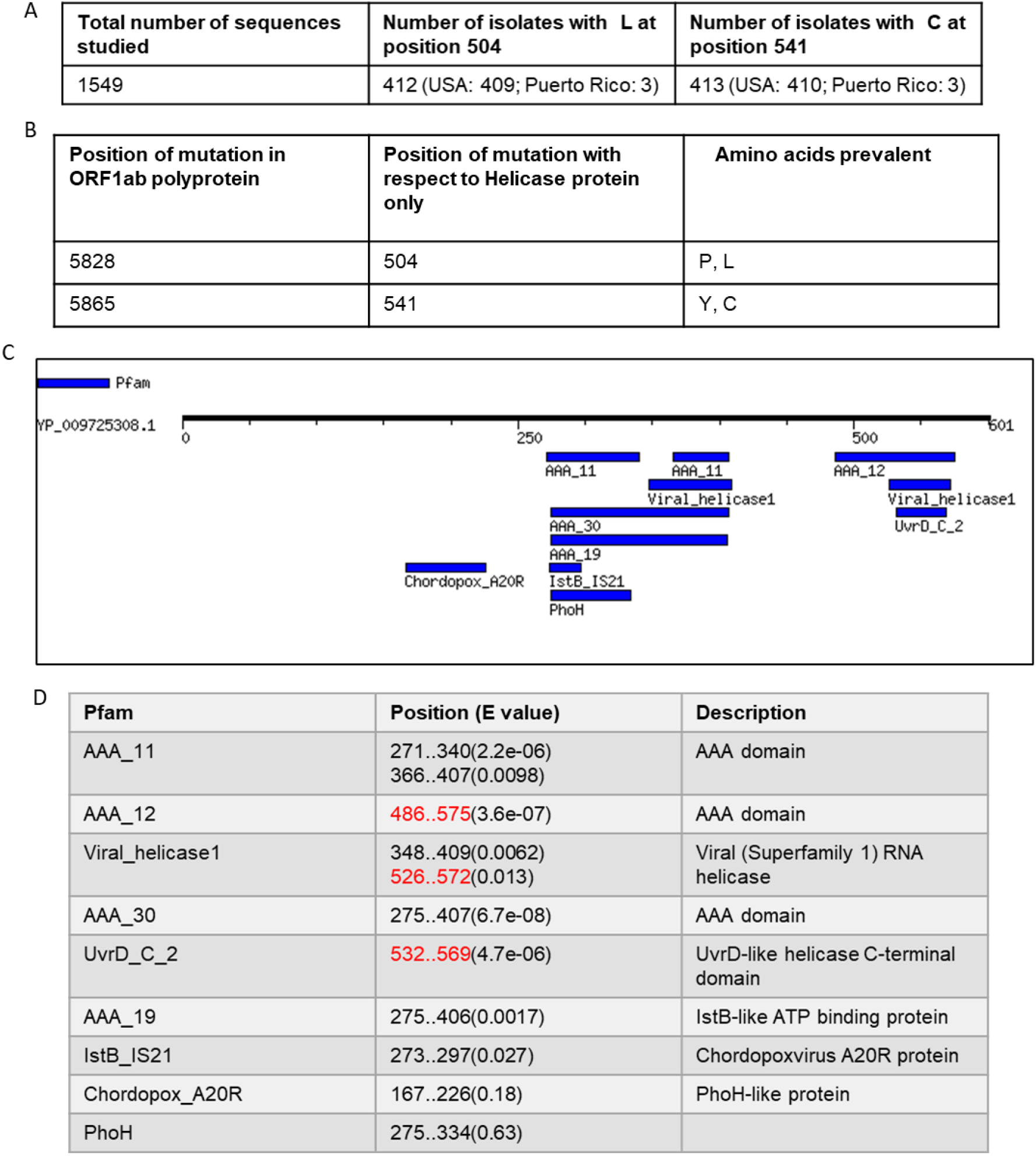
A, B. Mutations in SARS-CoV-2 helicase. C,D. Motif Scan of SARS-CoV2 helicase using MOTIF tool. Schematic diagram shows the motifs that the search engine identified and the same have been tabulated in the next panel. The positions of the mutations in the helicase are highlighted in red in the table.

The mutation P541C lie in the RNA helicase C terminal domain. Effect of this mutation on RNA binding and protein stability was studied and has been described later.

There is no crystal structure available to study structural biology of SARS-CoV2 helicase biology. Thus, to visualize these mutations in the context of 3D structure and future use for molecular docking studies, we generated homology models of P504L helicase and Y541C helicase using 6JYT.2 model available in SWISS MODEL repository as template. We selected the model/s based on maximum sequence similarity with the available template (Figure 2). These homology models could be used for detailed structure analyses, molecular docking studies and also target search in future by us as well as by others. Hence, these models will be deposited in public database for use. Here, we visualized the mutations in the context of full-length protein in 3D format using these structures. The mutations appeared to locally change the conformation of the helicase domain. Thus, we looked for alterations in the secondary structure of the protein using Chou and Fasman secondary structure prediction tool (Figure 3). In case of P504L, at the level of secondary structure there were loss of turns and introduction of helices and sheets as published earlier [5]. For mutation Y541C, a single helix was introduced (Figure 3).

**Figure 2.**
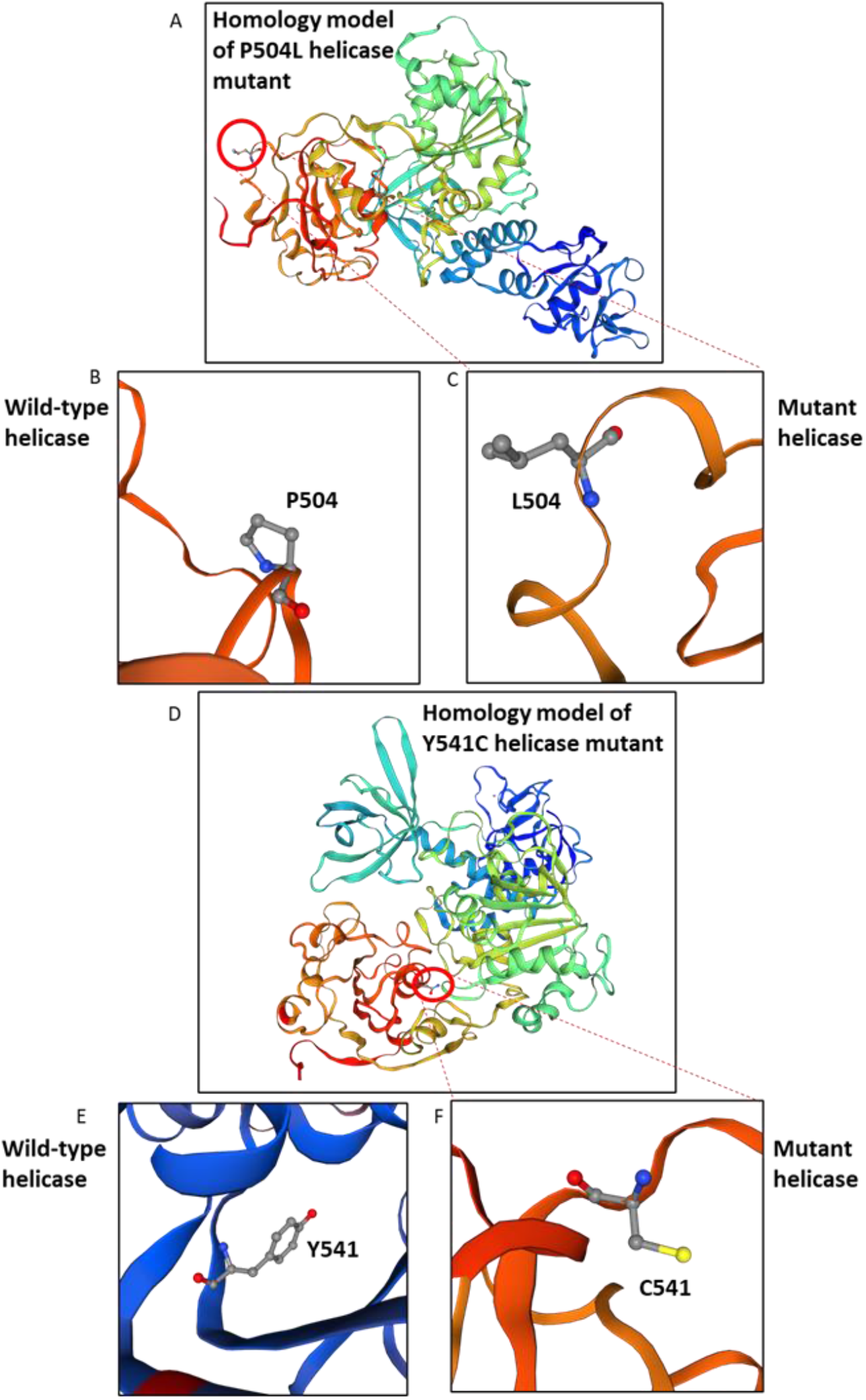
Homology Models of Mutant Helicases of SARS-CoV-2. A. Model of P504L mutant. B. Zoomed wild type helicase showing amino acid proline (P) at position 504. C. Zoomed P504L mutant helicase showing amino acid leucine (L) at position 504. D. Model of Y541C mutant. E. Zoomed wild type helicase showing amino acid tyrosine (Y) at position 541. F. Zoomed Y541C mutant helicase showing amino acid cysteine (C) at position 541.

**Figure 3.**
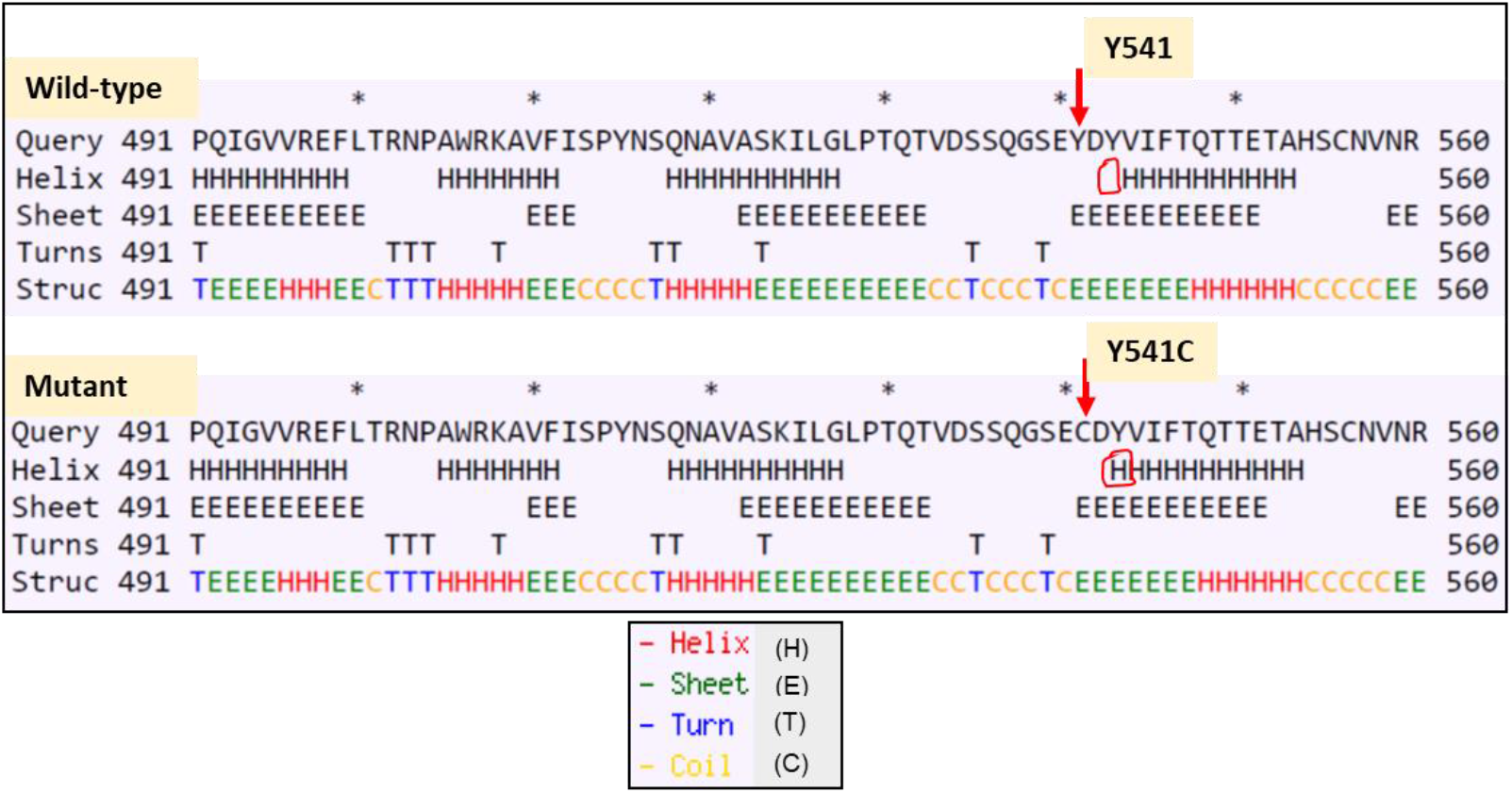
Effect of mutations on secondary structure of SARS-CoV-2 helicase. Alteration of secondary structure of helicase due to mutation Y541C. The secondary structure elements getting modulated have been marked with red boxes.

Changes in secondary structure can potentially alter stability of a protein. Thus, to study the function of mutant helicases, first we checked for the effect of mutations on the protein stability. Protein stability assessment using two independent prediction tools revealed similar results. Y541C mutation destabilized the helicase whereas P504L mutation could increase the stability of the protein. To correlate the structural modulation by these mutations with the function, we ran RNA binding analyses using mCSM NA server (Figure 4). RNA binding studies revealed that the replacement of tyrosine with cysteine decreased the affinity of the helicase protein for RNA template. Normal mode analysis/ protein dynamics study supported the mCSM data that the protein stability decreased and there was increase in protein flexibility for Y541C mutation (Figure 5). Atomic fluctuation and deformation energies complimented the observations. On the other hand, interestingly, mutation P504L increased both solvent accessibility and RNA binding affinity of SARS-CoV-2 helicase.

**Figure 4.**
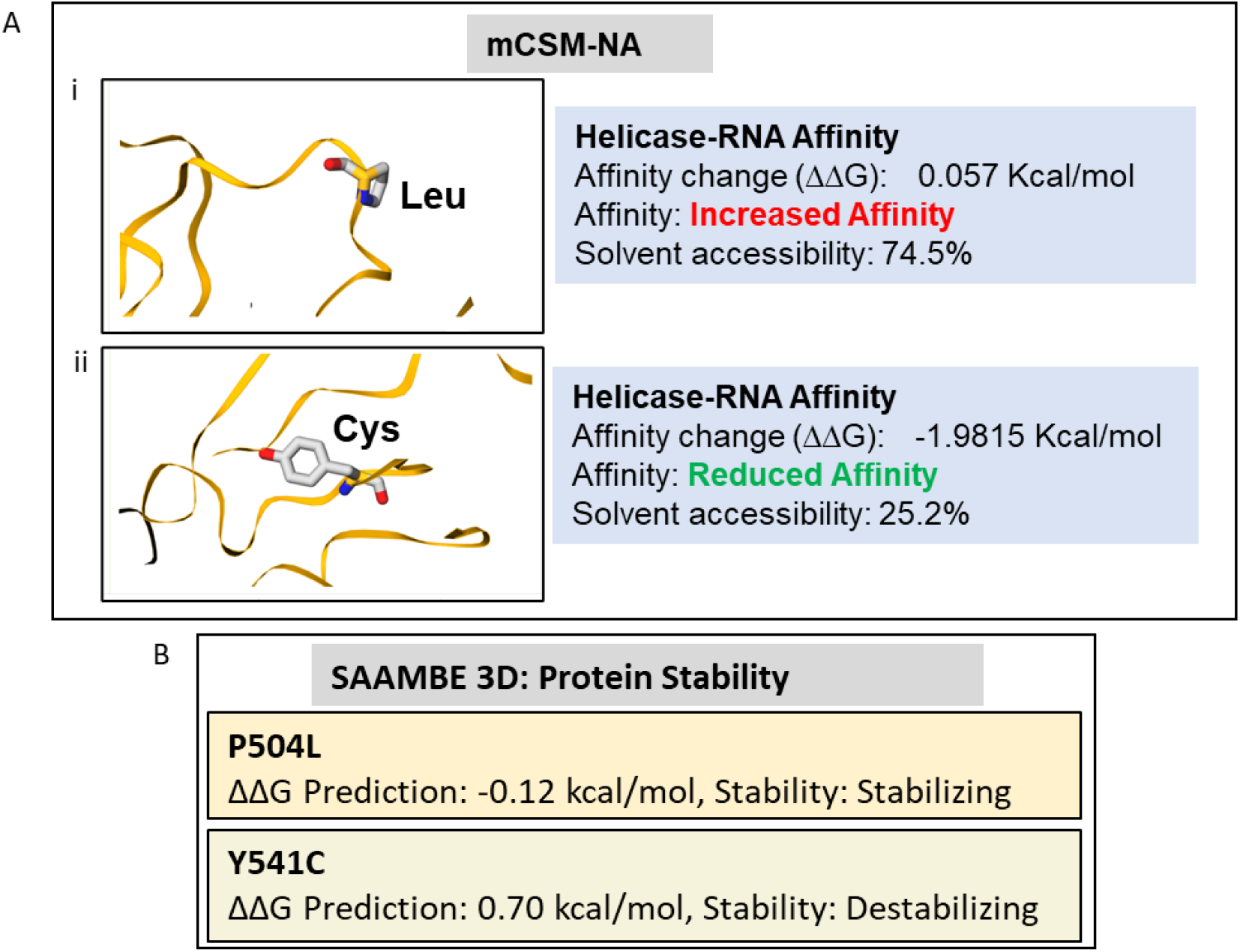
Effect of mutations on RNA binding ability and stability of SARS-CoV-2 helicase. A. mCSM analysis of RNA binding affinity of mutant SARS-CoV-2 helicases. B. SAAMBE 3D analysis to predict effect of mutations on stability of SARS-CoV-2 helicase. C. MUPro analysis to predict effect of mutations on stability of SARS-CoV-2 helicase structure.

**Figure 5.**
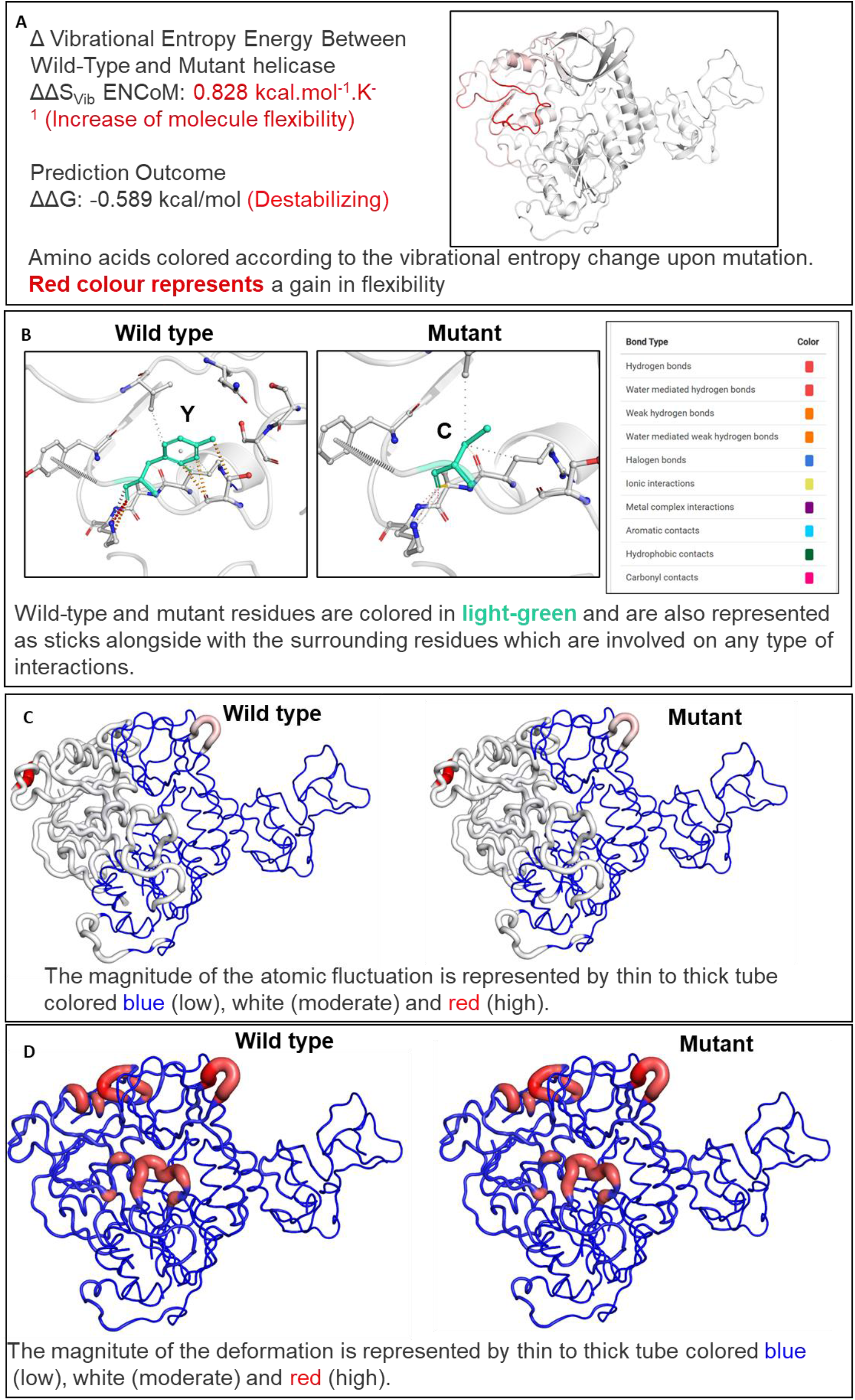
Effect of Y541C mutation on interatomic interactions and dynamics of SARS-CoV-2 helicase protein. A. Δ Vibrational Entropy Energy (Visual representation) B. Prediction of Interactomic Interactions C. Visual analysis of Atomic Fluctuation (amplitude of absolute atomic motion) D. Visual analysis of Deformation Energies (measure for amount of local flexibility)

Overall, this study revealed circulation of two mutations in the helicase protein of the SARS-CoV-2 viruses in a significant fraction of SARS-CoV-2 isolates. Our analyses indicated possible role of at least one of these mutations in enhancing affinity of helicase RNA interaction and thus replication. To further validate this model, these mutations could be incorporated in helicase protein and assayed for effect on RNA unwinding activity. Further, careful epidemiological surveillance and sequence analyses from more isolates would help understand the significance of these variants or lineages in evolution and spread of this virus. Such variants if found more virulent in nature could be specifically targeted focussing on the mutant helicase to design therapeutic candidates.

## Acknowledgements

We thank CSIR and AcSIR for support.

